# Evaluating the reliability, validity, and utility of overlapping networks: Implications for network theories of cognition

**DOI:** 10.1101/2021.09.28.462232

**Authors:** Savannah L. Cookson, Mark D’Esposito

## Abstract

Brain network definitions typically assume non- or minimal overlap, ignoring regions’ connections to multiple networks. However, new methods are emerging that emphasize network overlap. Here, we investigated the reliability and validity of one assignment method, the mixed membership algorithm, and explored its potential utility for identifying gaps in existing network models of cognition. We first assessed between-sample reliability of overlapping assignment with a split-half design; a bootstrapped Dice similarity analysis demonstrated good agreement between the networks from the two subgroups. Next, we assessed whether overlapping networks captured expected nonoverlapping topographies; overlapping networks captured portions of one to three nonoverlapping topographies, which aligned with canonical network definitions. Following this, a relative entropy analysis showed that a majority of regions participated in more than one network, as is seen biologically, and many regions did not show preferential connection to any one network. Finally, we explored overlapping network membership in regions of the Dual-Networks model of cognitive control, showing that almost every region was a member of multiple networks. Thus, the mixed membership algorithm produces consistent and biologically plausible networks, which presumably will allow for the development of more complete network models of cognition.

## 1. Introduction

Over the past 15 years, researchers have repeatedly demonstrated that brain regions can be consistently clustered together into a small number of spatially discontinuous groups based on the tendency for their activity during rest or performance of a task to be correlated in time. This measure of timeseries correlation between brain regions is referred to as functional connectivity (Biswal et al., 1995; Friston, 1994), and the groups of regions that demonstrate high mutual functional connectivity are referred to as intrinsic networks (Cole et al., 2013). Network neuroscience aims to identify these intrinsic networks in human functional magnetic resonance imaging (fMRI) data and understand how they interact in the service of behavior. Converging evidence from several different metrics indicates that intrinsic networks communicate through a relatively small subset of brain regions that are highly connected across multiple networks. For example, intrinsic networks exhibit many of the properties of a small-world architecture, including functionally segregated clusters with low numbers of cross-cluster connections and power-law distribution of degree (number of connections) across regions (Bassett & Bullmore, 2006). Likewise, graph theory metrics of participation coefficient and within-module degree, which assess the distribution of a region’s connections across networks, reveal a group of brain regions that are particularly strongly connected to multiple networks. Moreover, these properties underlie measures of network segregation and integration, such as modularity and between-network connectivity, which have been related to behavioral performance in a range of cognitive tasks (Baniqued et al., 2018; Cohen & D’Esposito, 2016; Kitzbichler et al., 2011; Mohr et al., 2016; Parkin et al., 2015).

While network neuroscience approaches have provided significant insight into the functional organization of the brain, global metrics like small-worldness, modularity, and between-network connectivity do not capture the role of individual brain regions in specific intrinsic networks. Even participation coefficient and within-module degree, which characterize region-level connectivity, only provide aggregate measures that collapse across networks.

Characterizing the specific networks to which individual regions connect would give further insight into functional organization of the brain that are lost with these measures. However, the methods implemented in most studies typically force non- or minimal overlap between the networks they use in their analyses or rely on previously published networks, almost all of which are likewise nonoverlapping (Glasser et al., 2016; Power et al., 2011; Yeo et al., 2011). The most common methods of network assignment rely on either clustering methods, which explicitly assign regions to a single network in a winner-takes-all fashion (Bassett & Bullmore, 2006), or independent components analysis (ICA), which tends to produce networks with minimal overlap in functional magnetic resonance imaging data as a matter of course (Langers, 2009). These methods of assignment likely lose important information about how particular brain regions may intrinsically connect to multiple networks.

However, methods exist that emphasize areas of overlap between networks by explicitly seeking out the meaningful connections between a region and all possible networks in the brain, which provides a straightforward and quantifiable way to capture how brain regions associate with multiple intrinsic networks. For example, Yeo and colleagues (2014) reported an early assessment of the differences between nonoverlapping and overlapping networks generated with ICA combined with a latent Dirichlet allocation algorithm (ICA-LDA). They found that overlapping networks largely captured the same topographies as nonoverlapping networks, indicating that they preserved known network structures. At the same time, the overlapping networks extended significantly beyond the boundaries of their nonoverlapping equivalents, revealing a wide range of pathways through which different networks could communicate. They also found areas of overlap that occurred entirely within the bounds of canonical networks rather than just at network boundaries, indicating that nonoverlapping assignment may obscure the areas that subserve this network communication. Critically, a large portion of the regions assigned to multiple networks were drawn from association areas, suggesting that these areas of overlap may be particularly important for coordinating between networks during higher-order cognitive processing. However, the assignments produced by this method only indicated whether or not a region was included in a given network. As such, while networks generated through ICA-LDA may be allowed to overlap, the method does not generate any additional metrics about each region’s association with each network, limiting its potential to expand our understanding of intrinsic networks.

More recently, Najafi and colleagues (2016) reported the first use of the mixed-membership algorithm (Gopalan and Blei, 2013) that generates novel information about multiple network membership above and beyond prior methods. This algorithm uses a stochastic block model to assign regions to a set of overlapping networks, producing an assignment vector for each region that indicates the strength of the region’s association with each network. This allows for intrinsic network overlap wherever the regions have a nonzero association with more than one network. Importantly, the area of overlap between two networks topographically indicates the regions that connect between those networks. This provides a direct map of which intrinsic networks each region contributes to and potentially to those regions that serve as communication bridges between networks. Najafi and colleagues applied the mixed membership algorithm to a set of fMRI data collected from 100 human subjects at rest and during several different tasks. Their analysis produced a set of overlapping networks that resembled networks produced using standard clustering methods, but extended into other areas of cortex as well. These results showed that multiple-network membership was common across a wide range of regions, including areas from all four lobes as well as subcortical structures.

The authors further demonstrated the within-sample reliability of these overlapping networks and explored the stability of the networks across dimensionalities (i.e., the number of networks specified for assignment). They also showed, for an arbitrary number of networks, good agreement between their overlapping networks and nonoverlapping networks generated with a common method in the same data. In addition, they further supported Yeo and colleagues’ (2014) previous conclusion that areas of overlap may support inter-network communication during cognition. Specifically, a region’s membership diversity, calculated as the Shannon entropy of each region’s assignment vector, was correlated with its functional diversity, or the number of cognitive domains it supported. Moreover, they identified a novel metric – “bridgeness” – that could separately identify regions that might serve as bottleneck gates and those that might support widespread integration.

Thus, methods such as the mixed membership algorithm give an intuitive map of regions’ multiple-network membership that is lost with previous methods, providing an important new tool for investigating and expanding network models of cognition. One prominent network theory of cognitive control, referred to as the Dual-Networks model of cognitive control (Dosenbach et al., 2007, 2008), posits the fronto-parietal (FP) and cingulo-opercular (CO) networks each supports distinct cognitive control processes. The FP and CO networks are composed of separate sets of regions and do not share any connections between them, which may be an artifact of the non-overlapping network methods used to define them. Using an overlapping method such as the mixed membership algorithm allows us to address answer this question, which has important implications regarding the specific mechanisms underlying network theories of cognitive control such as the Dual-Network model.

Here, we sought to replicate and extend Najafi and colleagues’ (2016) analyses of the reliability and validity of the mixed membership algorithm. We also aimed to further explore the utility of overlapping assignment for potentially refining network theories of cognition. We first tested the between-sample reliability of the mixed membership algorithm by calculating the topographical consistency of overlapping networks assigned from two different human subject samples collected under the same MRI protocols (Glasser et al., 2013; Van Essen et al., 2012). Next, we tested the method’s biological validity by calculating the topographical similarity of overlapping networks with nonoverlapping networks derived in the same data and relating these explicitly to well-established networks previously published in the literature. Finally, we tested whether the CO and FP networks have distinct sets of nodes, or whether some nodes in each of these networks are members of other networks. If the latter is true, revision of the Dual-Networks model, and likely other network models of cognition, will be necessary (Dosenbach et al., 2007, 2008).

## 2. Materials and methods

### 2.1. Datasets

The following analyses were performed on a 200-subject subset of the open access young adult data from the Human Connectome Project (S500 and S900 releases, Van Essen et al., 2012). Details on informed consent procedures and measures taken to ensure ethical and inclusive recruitment are described in detail by Van Essen and colleagues (2012, 2013). 100 subjects were selected as those having the lowest motion from the S500 release, calculated as the average framewise displacement across all four resting state scans. The second set of 100 subjects was selected at random from the remaining subjects from the S900 release after excluding the original 100 subjects. We selected 100 subjects for each subgroup as Najafi and colleagues (2016) previously demonstrated good internal reliability for this sample size, ensuring that each group in our split-half analysis would have reliable network definitions. The data used here included the de-identified preprocessed first LR acquisition resting state scans from each subject (1200 timepoints, TR = 0.72s, total scan time = 14 min 24s; Glasser et al., 2013). Each scan was further subjected to additional processing using functions available in the Analysis of Functional Neuroimages (AFNI) package (Cox, 1996). As has been previously reported (Hwang et al., 2017), the mean whole brain signal (-ort whole_brain_signal.1D) and frequencies outside of a band between 0.009 and 0.08 Hz (-band 0.009 0.08 -automask) were removed. Yeo and colleagues (2014) previously demonstrated similar results for overlapping assignment regardless of whether whole brain signal regression was used or not; we chose to apply whole brain signal regression in this analysis for its ability to remove artifacts due to motion and physiological signals and for its capacity to improve associations between resting state FC and behavior (Li et al, 2019). The analyses were conducted in volumetric space.

Illustrations of the networks are projected onto surface space for ease of whole-brain visualization.

### 2.3. Data processing

To prepare the data for network assignment analysis, the following preprocessing steps were followed using a combination of the AFNI package (Cox, 1996) and MATLAB software. A 1000-unit, 7-network derived version of the Global-Local parcellation (2mm resolution) by Schaefer and colleagues (2018) was used to define the regions of interest (ROIs) to be used for the correlation matrix. This parcellation provided the most fine-grained division of individual brain regions, especially in association areas such as the lateral frontal cortex.

The AFNI function 3dNetCorr (options: -fish_z, -in_rois) was used to extract region-wise, z-scored correlation matrices for analysis. ROIs with zero data were automatically removed from the correlation for each subject. Next, the data was imported into MATLAB and formatted for subsequent network assignment using a series of in-house scripts (available at https://github.com/savannahcookson/NetChar). Any ROIs excluded for a given subject were removed from all subjects, ensuring that all subjects had valid correlation values for all ROIs in subsequent analysis. As the mixed membership algorithm currently can only be applied to nonweighted, nondirectional networks, negative correlations were removed from the matrices; the the diagonal was converted to zeros to remove direct regional autocorrelations. The matrices were thresholded and binarized to retain only the top 10% of correlations to sparsify each subject’s correlation matrix; these were then averaged across subjects to weight the group-level correlation of a given pair of regions by the number of subjects with strong connections between those regions. The resulting average matrix was again thresholded to the top 10% of correlations to again sparsify the matrix and binarized for subsequent assignment with the mixed-membership algorithm (again, an unweighted assignment method).

For the split-half reliability analysis, we assessed the two subsamples of 100 subjects each separately; we will refer to the first and second subsamples as the “exploratory” and “confirmatory” datasets respectively for this report. The remaining analyses (validity and utility) were conducted on the combined 200 subjects from both subsamples to maximize the sample size used to define the overlapping networks in these analyses; we will refer this as the “combined” dataset.

The process of preparing the correlation matrices for network assignment and the actual assignment process both excluded several ROIs due to a lack of data in one or more participants. Correlation matrix processing excluded 25 ROIs in the exploratory dataset and 23 in the confirmatory dataset, where 21 were mutually excluded in both datasets. Overlapping network assignment excluded a further 36 ROIs in the exploratory dataset and 32 in the confirmatory dataset, with 29 mutually excluded. This resulted in a 939-region correlation matrix for the exploratory dataset and a 945-region matrix for the confirmatory dataset. These regions were mostly located in ventral frontal and anterior temporal cortex, and were likely due to signal dropout during data acquisition. Overlapping and non-overlapping network assignment from the combined dataset, and comparative analyses were restricted to those regions included in both the exploratory and confirmatory datasets, for a final count of 934 total ROIs.

### 2.4. Overlapping Network Assignment

Overlapping network assignment was conducted using the mixed-membership algorithm first reported by Gopalan and Blei (2013; package available at https://github.com/premgopalan/svinet) and applied to neuroimaging data by Najafi and colleagues (2016). Briefly, this program conducts overlapping network assignment by using an assortative stochastic block model to infer the probability (Bayesian estimation) that a given region is connected to each network, where the number of networks is pre-specified by the user. This produces an “assignment matrix” in which each region is given an association weight for each specified network.

We implemented this program on our data, specifying seven (7) networks for assignment and the number of ROIs included in the correlation matrix and setting the -linksampling option with the default threshold. Seven networks were specified to match the number of nonoverlapping intrinsic networks originally used to generate the regional parcellation used to define our ROIs as well as the number of networks reported in the previously published assignment results used to compare the networks in the current analysis to canonical networks from the literature. This permitted assessment of the extent of the differences in network topologies derived from overlapping and non-overlapping methods while keeping the base number of networks equivalent across all analyses. As the output of the mixed membership algorithm assigns each region a weighted association with each specified network, we binarized these assignment vectors without further thresholding (i.e., setting all nonzero values to 1, regardless of magnitude). This permitted direct comparison with the binary assignment topographies derived from nonoverlapping methods and allowed us to explore the broadest extent of potential overlap between networks.

### 2.5. Characterization of Assignment Consistency

Our first aim was to test the between-sample reliability of overlapping assignment by assessing mutual similarity between the overlapping network topographies produced by the mixed membership algorithm for the exploratory and confirmatory datasets, which were generated from separate subjects collected under the same protocol. To compare the overlapping networks from each dataset, we binarized each network without thresholding (i.e., all regions with a nonzero assignment value for that network were included in the topography) and then conducted a spatial Dice coefficient calculation via the AFNI command 3ddot (-dodice option, whole-brain mask) between each pair of networks between datasets.

To assess the significance of the Dice coefficients derived from this analysis, we conducted a 10,000-iteration permutation test. For each iteration, the assignment matrix region labels for the confirmatory dataset assignments were randomized and the Dice coefficients recalculated across the exploratory and shuffled confirmatory networks. Repeating this process for each iteration generated a null distribution of Dice coefficients for each pair of networks, which could be used to assess which of the Dice scores was significantly greater than chance. Significance was thresholded at □_FW_ = .05 family-wise corrected (Bonferroni procedure) across network pairs (49 comparisons total), for a within-comparison □ = .001. For visual comparison, we used these Dice estimates to estimate which of the seven networks from each set were the most mutually similar; i.e., to identify homologue networks between the samples. More specifically, we selected the seven pairs of networks between samples (without repetition) that maximized the total Dice score across network pairs. This process was also used to align other overlapping and nonoverlapping networks generated in later analyses. For completeness, we repeated this Dice similarity permutation testing between the overlapping networks generated for the combined dataset and each of those from the exploratory and confirmatory datasets to ensure that the networks used in subsequent analyses likewise captured the same topographies as either subsample.

### 2.6. Relation to Non-Overlapping Networks

Our second aim was to assess the validity of overlapping assignment by assessing the topographical similarities between overlapping and non-overlapping networks derived in the same data using methods commonly reported in the imaging literature. For example, Yeo and colleagues (2011) generated a set of non-overlapping networks in functional connectivity matrices derived from the resting state data of 1000 participants (500 participants each in a discovery and replication sample). They found stable clustering results at a resolution of 7 networks, and the topologies of these networks corresponded to previously reported functional networks. These networks are remarkably robust, having been replicated across a wide range of methodologies and datasets (Cole et al., 2014; Power et al., 2011). They have since become a mainstay of network neuroscience, commonly being used as a priori network definitions that are then used to assess integration and other metrics in other datasets.

Here, we used these extant network definitions (Yeo et al., 2011) to explore how overlapping networks related to the networks most commonly referenced in the literature. First, we applied a standard k-means clustering method to the averaged correlation matrices of the combined dataset here to define a set of non-overlapping intrinsic networks using standard techniques. First, we conducted the Dice coefficient permutation tests described above to compare the nonoverlapping networks to those defined previously by Yeo and colleagues (2011) to confirm their topographical consistency. Then, we likewise conducted a Dice coefficient permutation test to compare our non-overlapping topographies to the overlapping networks produced by the same data. In both of these permutation tests, the non-overlapping topographies identified by k-means assignment in the current dataset were shuffled to generate the Dice coefficient null distribution used for statistical testing.

### 2.7. Assessing Network Membership of Individual Regions

Our final aim was to perform an exploratory analysis to assess the overlapping network membership of regions canonically assigned to the FP and CO networks in the Dual-Networks model of cognitive control. To do this, we identified the regions that contained each of the 16 point-coordinates of the regions assigned to the FP and CO networks originally reported by Dosenbach and colleagues (2007, 2008) and extracted the assignment vectors of each region.

We also sought to characterize the extent these regions’ propensity for multiple-network membership relative to other regions in the brain. To do this, we calculated the total number of assigned networks and the entropy (*DKL*) of each region’s assignment vector relative to a uniform assignment reference distribution using the Kullback-Leibler divergence method from information theory. This method indexes the extent to which the actual assignment vector diverges from the reference distribution by estimating the amount of information about the assignment that would be lost by approximating it with that reference. In this method, regions that are assigned to more networks will have assignment vectors that more closely resemble a uniform distribution, resulting in a lower *DKL* value. We scaled these values to a range of zero to one for intuitive interpretation, such that regions with more uniform assignment tended toward zero (lowest divergence) and regions assigned to a single network had a value *DKL* = 1 (highest divergence). We plotted a histogram (100 bins) of these values across regions along with reference lines for the average *DKL* value as a function of the number of networks a given region was assigned to. We also included reference lines at the specific *DKL* values for each of the 16 FP and CO network regions.

## 3. Results

### 3.1 Comparison Between Samples

The reliability of the mixed membership algorithm was assessed by analyzing the consistency of the networks generated by two separate samples. Applying the mixed-membership assignment algorithm separately to the exploratory and confirmatory datasets produced two sets of seven networks that shared many similarities (Figure 1). Generally, assignment produced one network that primarily covered visual regions (Overlapping Network 4), two that covered mostly sensorimotor regions (Overlapping Networks 2 and 5), and four that were comprised of mostly association cortex.

**Figure 1.**
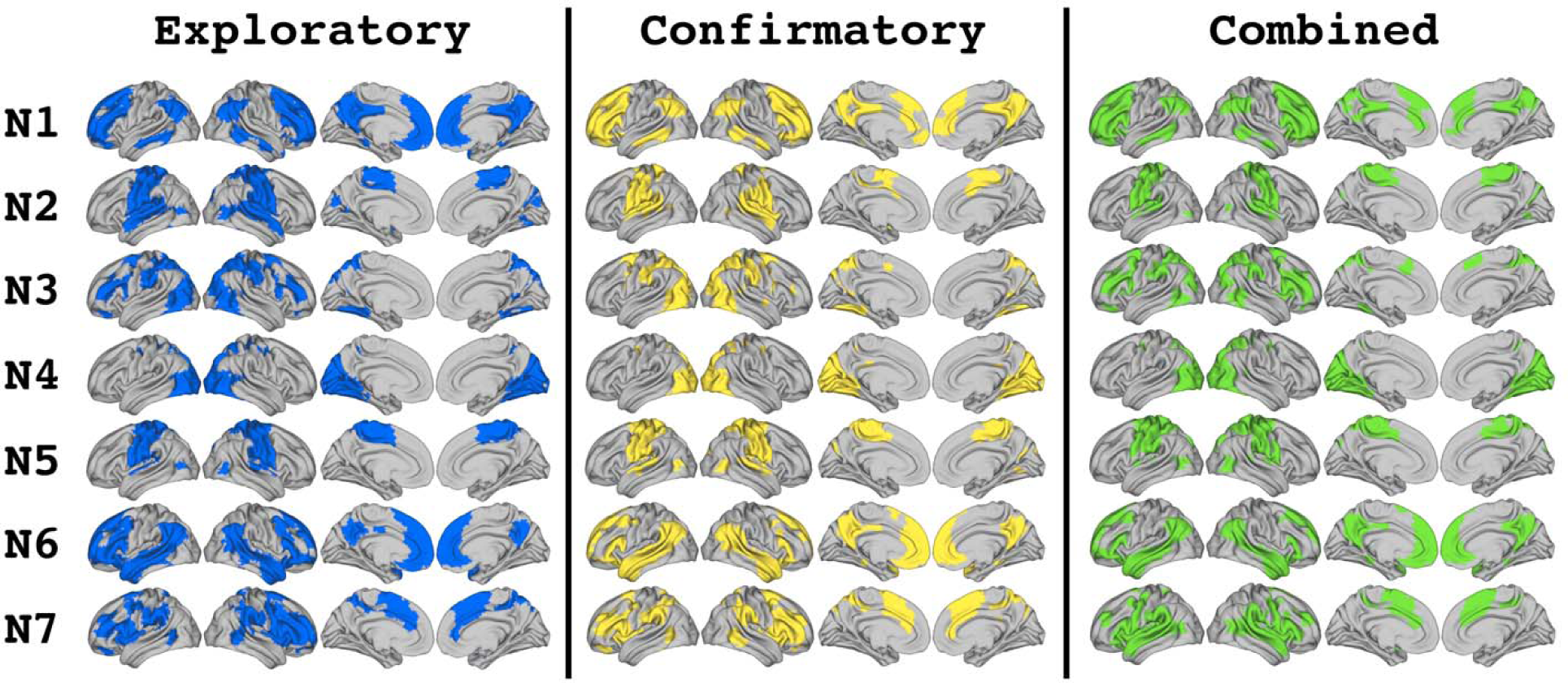
Overlapping network topographies generated by the mixed membership algorithm. Networks (N) are matched between sets to produce the largest overall similarity score, indexed by the sum of the Dice coefficients across network pairs.

The Dice coefficient analysis revealed that each network in one dataset was statistically similar to two to three networks in the other dataset (Table 1), consistent with a set of networks with significant areas of overlap between them. Homologous networks were defined by identifying seven mutually exclusive pairs of significantly similar networks between datasets while maximizing the sum of the Dice coefficients across pairs. These homologues showed strong between-pair similarity, with Dice coefficients greater than .75 for all seven pairs (mean: 0.86, range: [0.77:0.93]). There were also significant Dice coefficient similarities between non-homologue “secondary” pairs. These values were generally lower than those seen for the homologous pairs (mean: 0.56, range: [0.38:0.80]). Three of these secondary pairs were consistent between the exploratory and confirmatory networks. These included pairs between networks 1 and 6; networks 2 and 5; and networks 3 and 4. The remaining secondary pairs were: between Exploratory Overlapping Network (EON) 7 and Confirmatory Overlapping Networks (CONs) 1 and 2; and between CON7 and EON3.

**Table 1.** *Similarity between exploratory and confirmatory networks (Dice coefficient)*

|  | EON1 | EON2 | EON3 | EON4 | EON5 | EON6 | EON7 |
| --- | --- | --- | --- | --- | --- | --- | --- |
| CON1 | 0.89 | - | - | - | - | 0.69 | 0.43 |
| CON2 | - | 0.77 | - | - | 0.65 | - | 0.46 |
| CON3 | - | - | 0.85 | 0.50 | - | - | - |
| CON4 | - | - | 0.38 | 0.93 | - | - | - |
| CON5 | - | 0.80 | - | - | 0.91 | - | - |
| CON6 | 0.60 | - | - | - | - | 0.83 | - |
| CON7 | - | - | 0.50 | - | - | - | 0.87 |

*Note. Shows the dice coefficients of significantly similar pairs of networks between the exploratory and confirmatory datasets. "-" represents values that were not significant. EON: Exploratory overlapping network; CC: Confirmatory overlapping network.*

Comparing the overlapping networks from the exploratory and confirmatory datasets to those generated for the combined dataset (Figure 1) generally preserved this pattern of statistical similarity and homology for both sets of comparisons (Table 2). The patterns of similarity found between the combined dataset and each of the two comparison datasets were almost identical, with just one additional significant similarity score for the similarity between combined Overlapping Network (ON) 5 and EON7 that was not reflected in the scores for the combined and confirmatory networks. The patterns of both the combined-exploratory and combined-confirmatory similarities showed some additional similarities not reflected in the pattern of the direct comparison between the exploratory and confirmatory networks, likely because the networks derived for the combined dataset were partially informed by data from both of the split datasets. Overall, the mixed membership assignment produced a visually consistent set of networks with statistically verifiable homologues, supporting the reliability of this method for overlapping network assignment.

**Table 2.** Similarity between exploratory and confirmatory overlapping networks and overlapping networks from combined datasets (Dice coefficient)

|  |  | ON1 | ON2 | ON3 | ON4 | ON5 | ON6 | ON7 |
| --- | --- | --- | --- | --- | --- | --- | --- | --- |
| Exploratory | EON1 | 0.82 | - | - | - | - | 0.66 | - |
|  | EON2 | - | 0.73 | - | - | 0.67 | - | 0.54 |
|  | EON3 | - | - | 0.77 | 0.48 | 0.44 | - | - |
|  | EON4 | - | - | - | 0.95 | - | - | - |
|  | EON5 | - | 0.79 | - | - | 0.79 | - | 0.38 |
|  | EON6 | 0.66 | - | - | - | - | 0.81 | - |
|  | EON7 | 0.50 | - | 0.61 | - | 0.38 | - | 0.66 |
| Confirmatory | CON1 | 0.91 | - | - | - | - | 0.62 | - |
|  | CON2 | - | 0.64 | - | - | 0.51 | - | 0.63 |
|  | CON3 | - | - | 0.65 | 0.54 | 0.47 | - | - |
|  | CON4 | - | - | - | 0.92 | - | - | - |
|  | CON5 | - | 0.78 | - | - | 0.79 | - | 0.39 |
|  | CON6 | 0.53 | - | - | - | - | 0.91 | - |
|  | CON7 | 0.53 | - | 0.66 | - | - | - | 0.59 |

*Note. Shows the dice coefficients of significantly similar pairs of overlapping networks between the exploratory and confirmatory datasets and the overlapping networks produced for the combined datasets. "-" represents values that were not significant. EON: Exploratory overlapping network; CON: Confirmatory overlapping network; ON: overlapping network.*

### 3.2. Comparison with Non-Overlapping Networks

The validity of the overlapping networks generated by the mixed membership algorithm was assessed by comparing the overlapping topographies derived from the combined dataset to nonoverlapping networks in this same dataset. A set of seven nonoverlapping intrinsic networks was produced using a k-means analysis (Figure 2). We first compared these nonoverlapping networks to those previously reported by Yeo and colleagues (2011) to confirm that they represented similar topographies to those typically reported in the literature (Table 3). All of the nonoverlapping networks produced by k-means assignment were statistically similar to exactly one of the intrinsic networks described by Yeo and colleagues, with the exception of the “limbic” network. The HCP data used for the current analyses has weak signal in anterior temporal regions, many of which are included in the limbic network, so this result is not particularly surprising. In any case, the nonoverlapping networks defined in the current data generally aligned well with previously published intrinsic networks, insofar as the regions they contained were included in our data.

**Figure 2.**
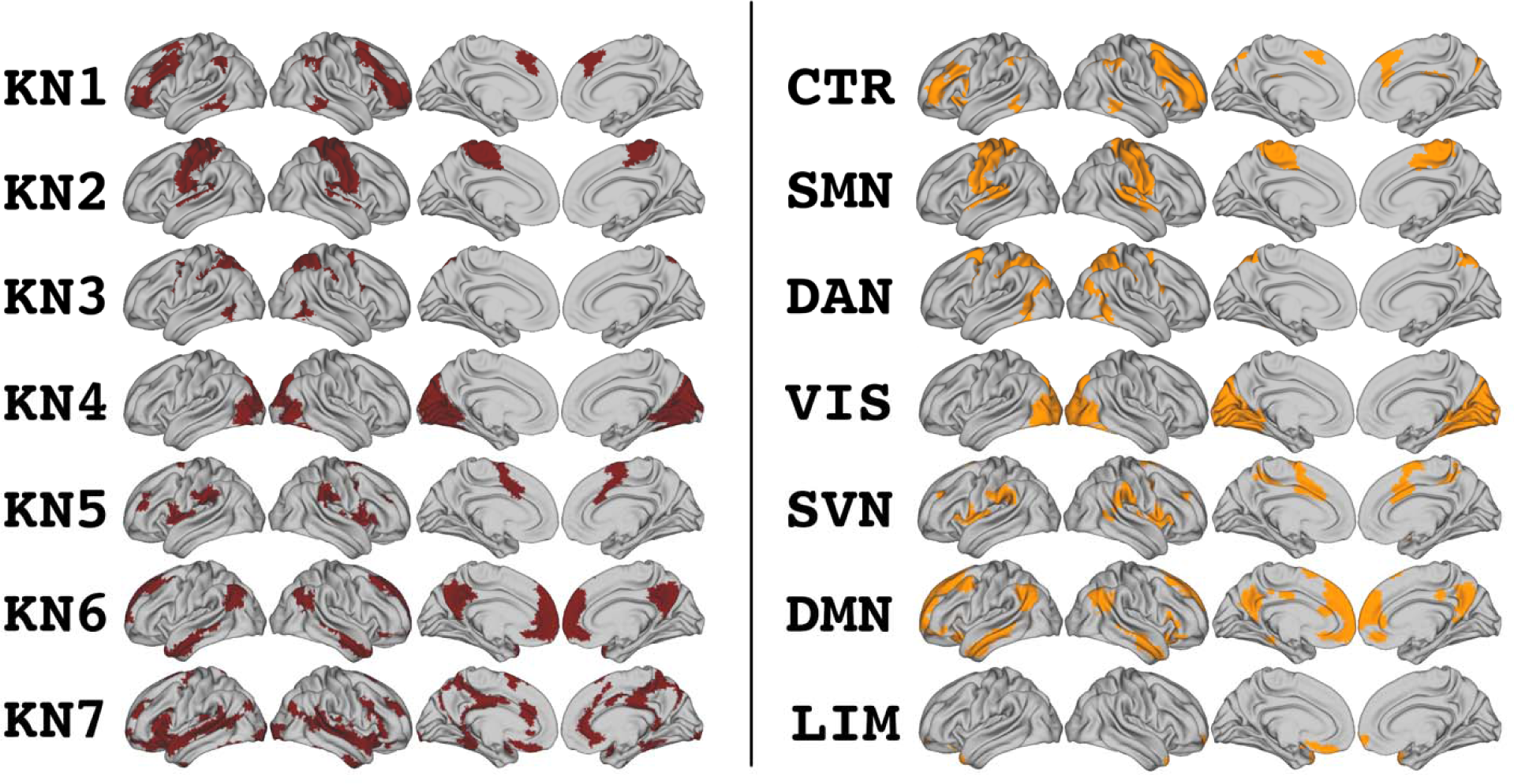
Topographies of networks generated by k-means clustering compared with original topographies reported by Yeo et al (2011). K-means networks (KN) for the current analysis are presented on the left; networks from Yeo et al are presented on the right with originally reported names. CTR: Control Network; SMN: Somatomotor Network; DAN: Dorsal Attention Network; VIS: Visual Network; SVN: Salience/Ventral Attention Network; DMN: Default Mode Network; LIM: Limbic Network. Brackets indicate the set of nonoverlapping networks whose topographies are included in each overlapping network.

**Figure 3.**
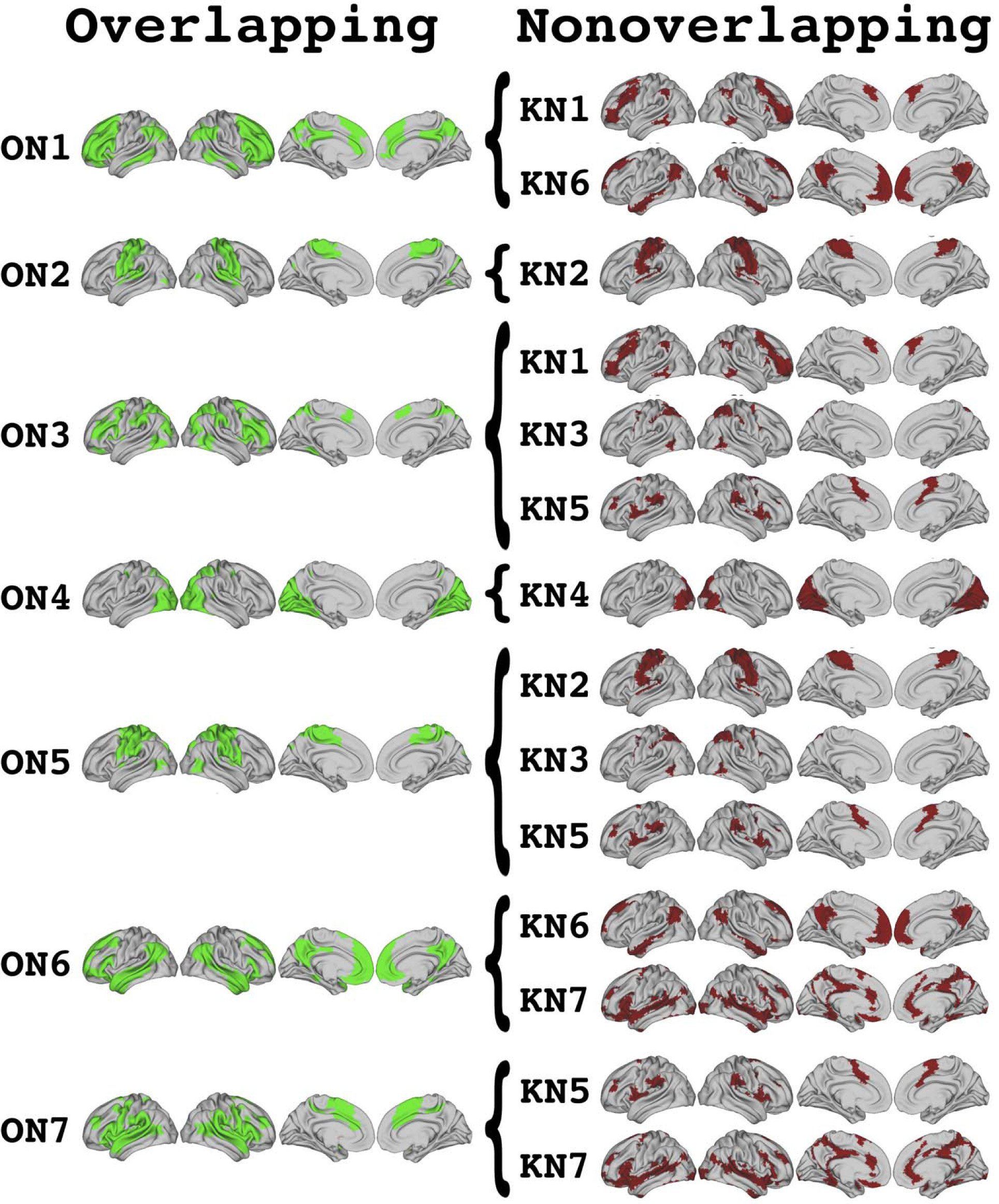
Visualization of which nonoverlapping networks are included in the topography of each overlapping network. Overlapping topographies are shown on the left in green (ON: overlapping network, combined dataset). Nonoverlapping networks (KN) are shown on the right in red.

**Table 3.** Similarity between k-Means networks and published networks (Dice coefficient)

|  | KN1 | KN2 | KN3 | KN4 | KN5 | KN6 | KN7 |
| --- | --- | --- | --- | --- | --- | --- | --- |
| CTR | 0.74 | - | - | - | - | - | - |
| SMN | - | 0.88 | - | - | - | - | - |
| DAN | - | - | 0.71 | - | - | - | - |
| VIS | - | - | - | 0.90 | - | - | - |
| SVN | - | - | - | - | 0.77 | - | - |
| DMN | - | - | - | - | - | 0.73 | - |
| LIM | - | - | - | - | - | - | - |

*Note. Shows the dice coefficients of significantly similar pairs of networks between k-means networks from the current analysis and networks previously reported by Yeo and colleagues (2011). "-" represents values that were not significant. KN: k-Means network; SVN: Salience/Ventral Attention Network; SMN: Sensorimotor Network; CTR: Control Network; DAN: Dorsal Attention Network; DMN: Default Mode Network; VIS: Visual Network; LIM: Limbic Network.*

We next compared the topographies of the nonoverlapping networks from the combined dataset to the overlapping networks produced in the same dataset using the mixed membership algorithm. In this analysis, the overlapping networks were significantly similar to one to three nonoverlapping networks, indicating that each overlapping network represented one or a combination of a small subset of nonoverlapping networks (Table 4; Figure 4). Two overlapping networks were similar to exactly one non-overlapping network; ON4 aligned with the “visual” network, and ON2 with the “somatomotor” network. The remaining five overlapping networks were similar to two or three non-overlapping networks. ON2 and ON4 showed relatively similar dice coefficients with their respective KN homologues as were seen for network homologues in our split-half and published network analyses. On the other hand, the dice coefficients for the remaining network homologues were lower than in other analyses, indicating that there was not a one-to-one mapping of overlapping networks to canonical intrinsic topographies. Instead, each overlapping network appeared to combine large areas of two or three nonoverlapping topographies.

**Figure 4.**
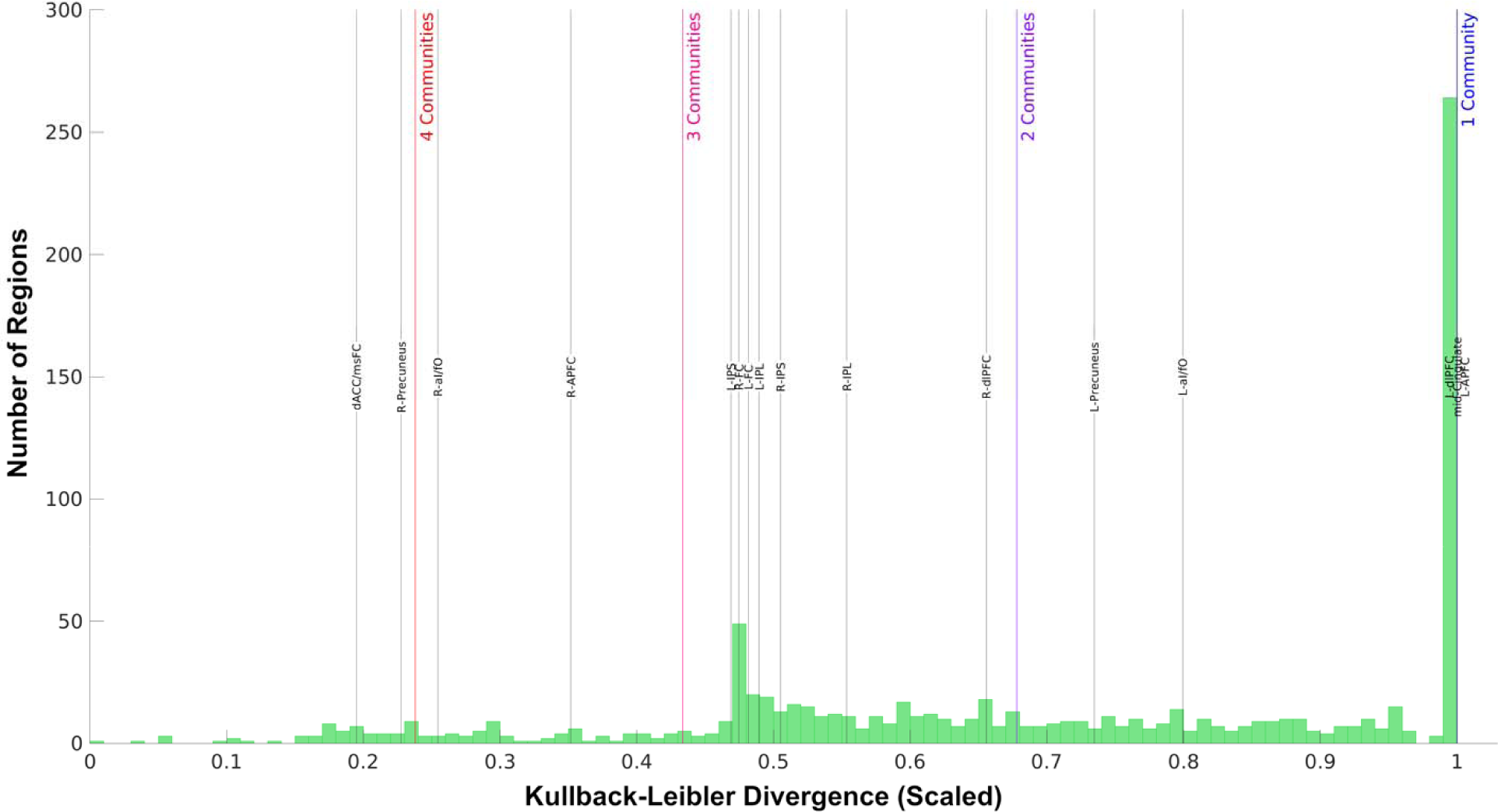
Distribution of relative entropies across brain regions. Histogram of the relative entropies across the 937 regions included in overlapping network assignment. Colored vertical bars represent the average relative entropy for all regions assigned to the labeled number of networks. Black vertical bars show the specific relative entropies for the 16 regions of the FP and CO networks.

**Table 4.** Similarity between k-Means networks and overlapping networks (Dice coefficient)

|  | ON1 | ON2 | ON3 | ON4 | ON5 | ON6 | ON7 |
| --- | --- | --- | --- | --- | --- | --- | --- |
| KN1 | 0.54 | - | 0.35 | - | - | - | - |
| KN2 | - | 0.83 | - | - | 0.57 | - | - |
| KN3 | - | - | 0.42 | - | 0.35 | - | - |
| KN4 | - | - | - | 0.73 | - | - | - |
| KN5 | - | - | 0.26 | - | 0.21 | - | 0.44 |
| KN6 | 0.34 | - | - | - | - | 0.65 | - |
| KN7 | - | - | - | - | - | 0.44 | 0.44 |

*Note.* Shows the dice coefficients of significantly similar pairs of networks between the k-means networks and overlapping networks generated for the combined datasets. "-" represents values that were not significant. KN: k-Means network; ON: overlapping network.

### 3.3. Exploring Network Models with Overlapping Assignment

In this final analysis, we examined the extent of multiple-network membership in regions across the brain and explored whether the regions of the FP and CO networks, which were previously defined using a nonoverlapping method, were members of other networks. Across the brain, the weighted overlapping network assignments for the combined dataset were extracted for each region and subjected to a relative entropy analysis. This analysis revealed a large number of regions (263 out of 937) with entropy values at 1, with the rest of the brain regions with entropy values less than 1 distributed relatively evenly across the full range of values (*DKL* range: [0:0.9912]; Figure 4). While many of regions were assigned to a single overlapping network (i.e., *DKL* approaching 1), many more (i.e., DKL < 1) were assigned to multiple networks, with a maximum of four assigned networks for some regions (Figure 4).

Next, we extracted the assignment vectors for the brain regions of the FP and CO networks (as defined by a non-overlapping assignment method) that form the basis for the Dual Networks model of cognitive control. These regions were members of all seven of the overlapping networks, where the most common assignments were to networks 1, 3, and 7 (Table 5). ON3 visually appeared most like the canonical FP network, further incorporating regions typically associated with the Dorsal and Ventral Attention networks; ON7 likewise was primarily composed of regions associated with the canonical CO (or Salience/Ventral Attention network from Yeo and colleagues, 2011) network with additional areas from superior temporal cortex. Accordingly, most regions assigned to ON3 were nodes of the FP network, while most regions assigned to ON7 were nodes of the CO network; however, both of these overlapping networks contained regions from both the FP and CO networks. ON1 on the other hand did not resemble any one particular nonoverlapping network, instead combining regions from the FP and Default Mode networks in one topography. ON1 showed broad inclusion of regions from both the FP and CO networks.

**Table 5.**
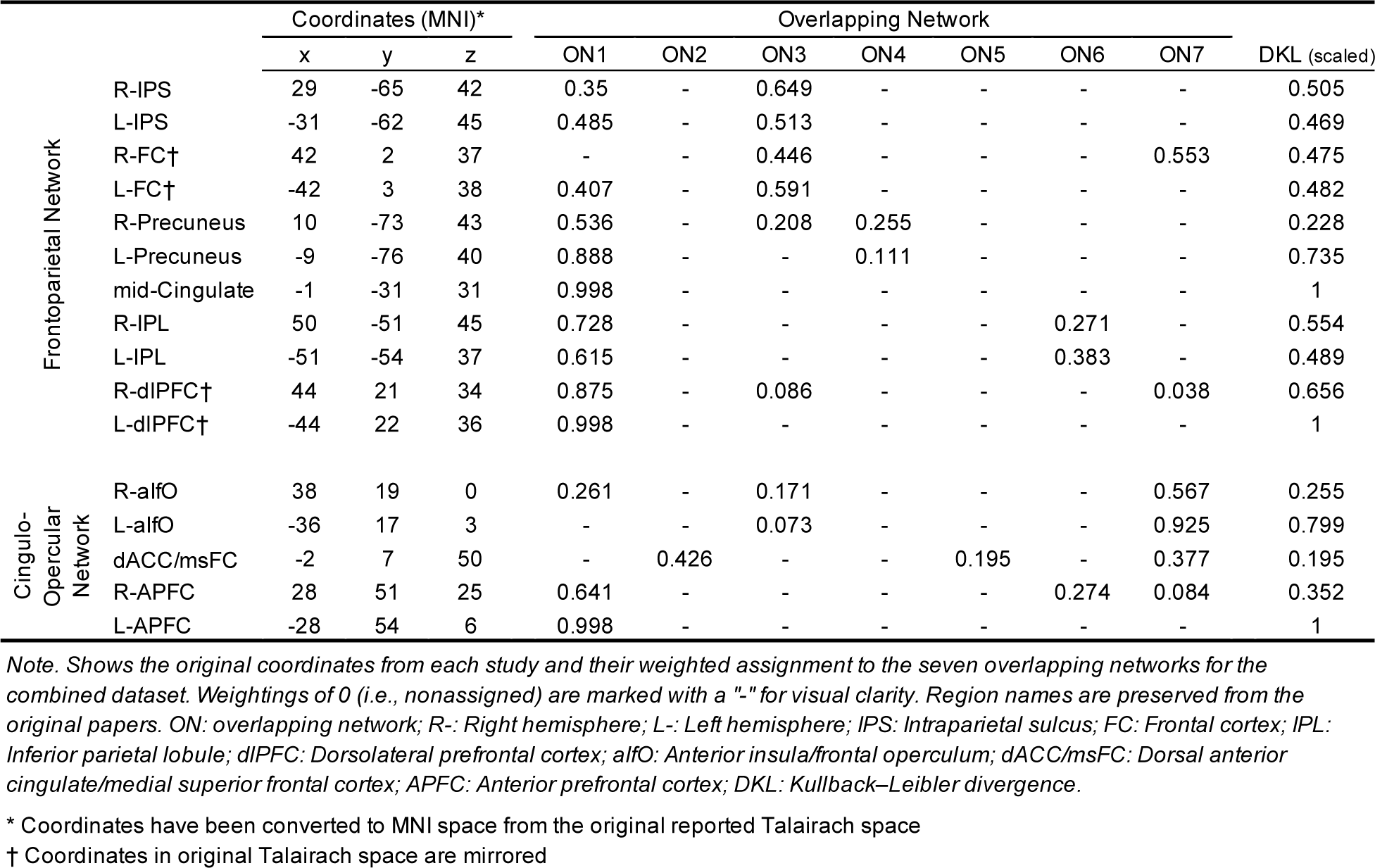
Overlapping network membership and graph theory properties of LFC regions

Most regions of the FP and CO networks were assigned to two or three overlapping networks, with only the left anterior prefrontal cortex (L-APFC) and mid-cingulate regions assigned to a single network. Accordingly, most of regions from the FP and CO networks had relative entropy values much less than one. The regions with the lowest values – the dorsal anterior cingulate/medial superior frontal cortex (dACC/msFC), right precuneus, and right anterior insula/frontal operculum (R-aIfO) – were the three regions assigned to three overlapping networks, the highest number of networks assigned for any FP or CO network regions.

## 4. Discussion

The results of our analyses indicate that the overlapping networks generated by mixed-membership assignment have good between-sample reliability, and thus produce consistent overlapping networks across subjects. They also meaningfully recapitulate structures seen in nonoverlapping networks both generated in the same data and published in the greater literature, indicating that the mixed membership algorithm produces equally valid network assignments as previous assignment methods. At the same time, the overlapping networks revealed relationships between subgroups of nonoverlapping topographies that are obscured with nonoverlapping definitions, indicating that they may better capture the biological reality of the brain’s network structure. Finally, these results indicate the utility of the mixed membership algorithm for revealing regions’ involvement in networks that are obscured by nonoverlapping methods. Together, these results demonstrate that overlapping network assignment is a practical and accessible tool. Moreover, they highlight the pressing need to consider how network overlap can be accounted for in our existing network models of behavior.

### 4.1. Reliability of overlapping network assignment

Overall, the overlapping networks generated by the mixed-membership algorithm showed good agreement between two samples based on the Dice coefficient permutation test. All of the networks derived from the exploratory dataset had significant similarity with two or three networks from the confirmatory dataset. Homologous network pairs could be identified for all seven identified networks, along with additional similarities that reflected consistent areas of overlap between networks. Moreover, these patterns were replicated when comparing each of the sets of networks from the split-half datasets to those generated across the full combined dataset. Together, these findings suggests that the mixed-membership assignment method produces reliable networks between samples.

Najafi and colleagues (2016), who originally reported the use of the mixed-membership algorithm on brain imaging data, previously demonstrated that the mixed membership algorithm is a reliable method within sample using a bootstrapped reliability analysis. The current results extend those findings to now account for between-sample reliability as well. Additionally, Najafi and colleagues conducted their analyses in surface space whereas the current results are reported in volumetric space. The results from both spaces show similar reliability and validity, demonstrating the mixed-membership algorithm’s robustness in both spaces. Notably, direct comparison of the networks produced here and those found by Najafi and colleagues is not possible due to differences in the number of networks specified for analysis (Najafi et al: 6; here: 7). In their original report, Najafi and colleagues conducted a stability analysis across network dimensionality, which revealed a slight peak at six networks. However, the stability level was similar for seven networks in that analysis, and the use of seven networks here permitted us to relate our overlapping networks to previously reported network definitions. Nonetheless, there were marked qualitative similarities in the spatial topographies of the overlapping networks reported by Najafi and colleagues and the current study. Future research is necessary to explore how the number of networks used for assignment might lead to comparable and/or contrasting topographies and the best resolution for overlapping assignment for various applications.

An additional difference between the current research and the study by Najafi and colleagues is the computational demand of the methods employed for network assignment. Najafi and colleagues applied an iterative assignment process that aligned the most similar network topographies across iterations and generated a consensus assignment that accounted for variations in assignment region-by-region. Here, we attained similar reliability across networks between samples with a single round of assignment, lowering the computational demands of the assignment significantly. Notably, the network assignments produced by the mixed membership algorithm were reliable between samples even though the assignment weights were ignored in the Dice coefficient analysis, further indicating their overall robustness. We have made our processing scripts available as an easily accessible open-source download (https://github.com/savannahcookson/NetChar), which should facilitate more widespread adoption of the mixed membership algorithm for network assignment in future studies of brain networks.

There were some small qualitative differences in the network topographies derived from the exploratory and confirmatory datasets, most notably in the areas of medial frontal and mid-prefrontal cortices, superior temporal cortex, and parietal cortex, which might call into question the overall reliability of mixed membership assignment. However, given the otherwise high consistency in network assignment across the two datasets, this seems unlikely. Instead, this inconsistency may reflect particularly high inter-subject variability. Recent work identifying “network variants” across individuals using non-overlapping methods may be a source of this variability in network assignment across datasets. Gordon and colleagues (2017) first reported these network variants in a set of ten subjects with a large amount of high-quality individual data. Generating a non-overlapping network parcellation for each subject individually revealed variations in network assignment in certain areas that were not present in the group average.

These areas included anterior mid/inferior and ventromedial prefrontal cortices, middle cingulate cortex, and superior parietal cortex, which are similar in location to the areas that were inconsistently assigned here. In a follow-up study using the same dataset, Gratton and colleagues (2018) demonstrated that these variants were stable individual features across task demands, indicating that these variants were likely intrinsic. Moreover, Seitzman and colleagues (2019) demonstrated that these variants are trait-like, suggesting that they are stable over time within individuals. They also showed that participants could be divided into groups that show similar variant structures across individuals. Future research using the mixed-membership algorithm will provide an opportunity to further our understanding of these network-variants and their influence on group-level network assignment.

### 4.2. Validity of overlapping network assignment

To determine the relationship between overlapping and non-overlapping networks, we applied a k-means clustering algorithm to the correlation matrix for our combined datasets to produce seven non-overlapping networks. The nonoverlapping networks we identified were generally topographically consistent with networks previously reported by Yeo and colleagues (2011). Moreover, the overlapping networks we identified showed meaningful recapitulation of topographic relationships that would be expected from nonoverlapping assignment while appearing to combine subgroups of those nonoverlapping topographies in specific ways. Two overlapping networks showed a strong similarity to one single nonoverlapping network, one visual and one somatomotor; the remainder were composed of combinations of multiple networks comprised of brain regions in association areas. Najafi and colleagues (2016) also performed a comparative analysis between overlapping and non-overlapping networks in the same dataset by calculating the spatial correlation of their topographies, with similar conclusions. Four of their six overlapping networks had a single nonoverlapping homologue, two of which (communities 1 and 2 in their original report) included visual and somatomotor cortices respectively as seen in our results here. The other two networks were composed of association cortex from widespread frontoparietal and temporal regions.

Given that all seven of our overlapping networks have topographic consistencies with non-overlapping networks that match those from the literature, it seems unlikely that overlapping assignment is capturing fundamentally different or novel networks from those identified using non-overlapping assignment to define the networks. Instead, the overlapping networks appear to replicate the nonoverlapping topographies either individually or in limited combinations.

Altogether then, overlapping and nonoverlapping assignment methods are capturing similar mathematical patterns in the functional connectivity data. It is tempting then to ask which method better captures the “ground truth” of the brain’s network topographies; however, this begs the question of how the ground truth can be achieved. Shinn and colleagues (2017) point out that there is no epistemological reason to believe that there is a community structure in a given network a priori, and further that there is no possible method to determine the ground truth of the correct resolution or decomposition of that community structure. Instead, a better question is to ask is which method is more biologically plausible.

First, there is evidence that individual brain regions are anatomically connected to multiple networks, not just regions within their own network. For example, seminal work using histological tracing in primates (for review, see Goldman-Rakic, 1988) demonstrated that frontal region 46A and parietal region 7A, regions assigned the FP network in modern analyses, are directly connected to the superior temporal sulcus (e.g., a region assigned to the default mode network), post-cingulate gyrus (e.g., a region assigned to the salience/ventral attention network), and area 19/peristriate cortex (e.g., a region assigned to the visual network). Work in humans indicates that functional connectivity captures not only these direct connections but also many indirect connections (Honey et al., 2009), suggesting that network overlap is even broader in functional data. Accordingly, evidence from human brain imaging studies demonstrates that regions show flexibility in network affiliation, both at rest and during task performance (Cole et al., 2014; Khambhati et al., 2018; Pedersen et al., 2018). Moreover, studies using other techniques that have allowed for some level of network overlap (Bijsterbosch et al., 2019; Yeo et al., 2014) likewise have demonstrated that regions can be assigned to multiple networks. Finally, many studies have highlighted the behavioral importance of “hub” regions that share a high number of connections with regions outside of their assigned network (Cole et al., 2013; Hwang et al., 2017; Power et al., 2013). Ultimately, there is no distinction between which network(s) a region is a “member” of and which they merely “connect” to, a point previously made in Pessoa’s (2014) recent review of brain networks; these hubs are simply members of multiple networks. While the connections of these hubs may be able to be ascertained with nonoverlapping methods, though, there may be other regions that are not identified as hubs that may nonetheless be members of multiple networks. These regions’ involvement in multiple networks is effectively lost with nonoverlapping assignment.

Thus, the existing literature broadly indicates that overlapping networks do indeed better capture the brain’s large-scale network organization. This notion is consistent with studies of task-related networks. For example, Nee (2021) demonstrated a rostro-caudal gradient of network membership related to the processing of three distinct cognitive control factors, with extensive overlap between adjacent networks. Each of the three networks included regions from multiple intrinsic nonoverlapping networks. More specifically, each task-related network included combinations of regions from the FP network, default mode network, salience network, and dorsal attention network. Notably, the combined topography of these four task-related networks is similar to the topography of one of the overlapping networks we identified (i.e., ON1). Future research should relate these and other task-related networks to overlapping network topographies to better understand how network overlapping drive dynamic network reorganization during task performance. More importantly, it is vital that network neuroscience embrace methods that explicitly adopt an overlapping network perspective to support more biologically plausible interpretations of the brain’s network organization and avoid losing information about which regions connect to multiple networks.

### 4.3. Utility of overlapping network assignment

A unique feature of the mixed membership algorithm, like many overlapping assignment methods, is that it generates an assignment vector for each region, which captures the probability that that region is a member of a particular network. At a whole-brain level, we assessed which overlapping networks each region was assigned to. Combined with our analysis of the similarity between our overlapping and nonoverlapping networks, this gave us an intuitive map of how regions in different nonoverlapping networks shared membership across multiple networks (Figure 5). An analysis of the relative entropy of assignment for each of these regions revealed a broad spectrum of connectivity distributions across different overlapping networks.

**Figure 5.**
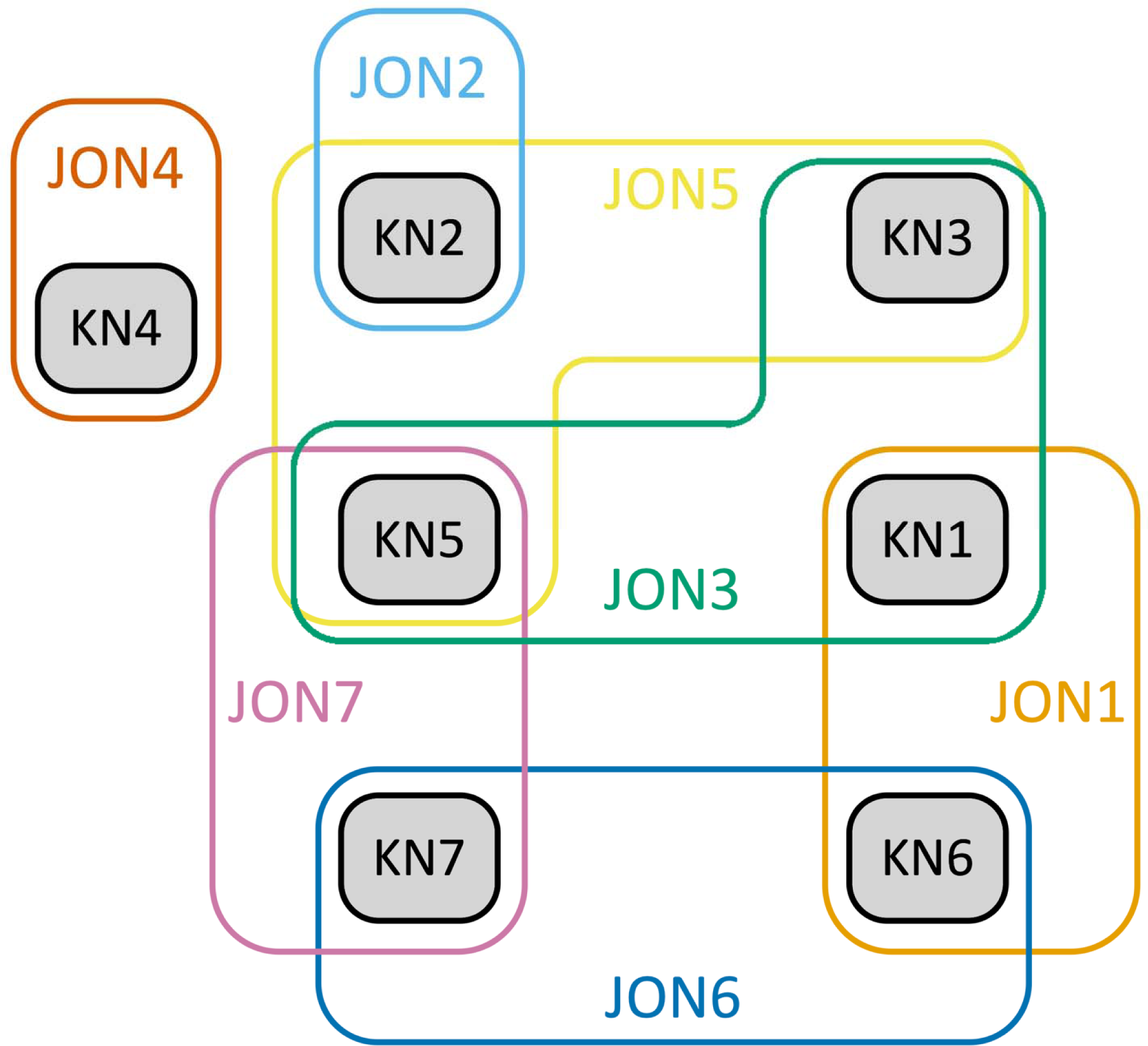
Visualization of nonoverlapping network topographies that significantly make up each overlapping network. K-Means networks (KN) are presented in black. Each Overlapping Network (ON, combined dataset) is then presented in a different color encircling the set of k-means topographies that is significantly represented in the overlapping network.

That is, regions did not universally show preferential assignment to one network over another; instead, many regions showed relatively even assignment to two or even more networks simultaneously. This yet further supports the need to move toward using overlapping methods that do not obscure this information.

The goal of any network model of behavior is to capture which networks support that behavior and how they interact with one another. A common method of exploring brain network interactions is to assess their integration and/or segregation during different tasks (see Sporns, 2013 for review) by calculating by assessing the connectivity between regions in one network and regions in another network and comparing this value between task conditions or between task and rest. Previous research using this method has demonstrated that the FP and CO networks demonstrate increased integration during performance of a task requiring the integration of FP- and CO-related processing relative to a simple categorization task (Cohen et al., 2014), which is taken as evidence that these two networks indeed interact during cognitive control. Similarly, Menon (2011) has proposed a “Tri-Network” model of psychopathology that proposes that clinical symptoms manifest as a function of disorderly interactions between the FP network, the default mode network, and the salience network.

However, any speculation about interactions between networks in models such at the Dual-Networks model of cognitive control or the Tri-Network model of psychopathology is incomplete without accounting for regions that are members of multiple networks. To illustrate, we extracted the assignment vectors for each of the regions of the FP and CO networks to determine their possible assignment to all of the overlapping networks that we identified.

Collectively these regions were members of all seven networks, indicating the potential involvement of more networks than just the FP and CO networks in the service of cognitive control. Even the relatively restricted set of FP regions analyzed here were members of five networks when networks were allowed to overlap, suggesting that any model involving the FP network must account for interactions with many other networks. Moreover, our results revealed a novel network combining regions from the FP and Default Mode networks that is not found with nonoverlapping methods; this and other potential “hidden” networks will need to be accounted for as well. There is indeed mounting evidence that several additional networks are involved in various aspects of cognitive control, including the dorsal attention network (Badre & Nee, 2018; Ito et al., 2017; Nee, 2021), salience/ventral attention network (Seeley et al., 2007), and even default mode network (Smith, Mitchell, & Duncan, 2018; see also Cocchi, Zalesky, Fornito, & Mattingley, 2013). Thus, a complete network model of cognitive control needs to be updated to account for the role of these additional networks and the complex interactions that occur as a function of their overlapping nature.

### 4.4. Limitations

As the use of the mixed-membership algorithm, and indeed the application of overlapping networks more generally is nascent in network neuroscience, there are several limitations in the current analyses that will need to be addressed and clarified in future work. As with all network analyses, the current study conducted several preprocessing steps to prepare the correlation matrices for assignment. These methods include typical steps of zeroing negative correlations, correlation thresholding, and edge binarization. While these are commonly used across studies in the network neuroscience literature, they are primarily (as in the current study) employed as a matter of mathematical convenience or necessity. Additionally, the mixed membership algorithm has as yet only been implemented for unweighted, undirected, and sparse correlation matrices. While graph theoretical methods for network assignment of directed, weighted, dense correlation matrices no doubt exist, versions that use stochastic block models or other forms of overlapping assignment have not yet been reported in the neuroscience literature. A related issue is global signal regression, as was used in the data analyzed here. Global signal regression may improve specificity of positive correlations, but also can induce spurious anti-correlations in functional connectivity analyses. It will be necessary to characterize the impact of these preprocessing steps and parameters on the final results of overlapping assignment in future research.

Finally, the HCP dataset used for these analyses collected a significant number of samples from genetic twins. As the subsets of subjects selected for this analysis were randomized, they may have included multiple samples from these twin groups. It is possible that inclusion of twins may have artificially inflated the reliability estimates between samples if one twin was included in each subgroup in the split-half analysis. At the same time, it is equally likely that both twins from a group were included in the same subgroup, which could have resulted in lower overall reliability by biasing one subgroup toward an overly represented network structure in the data. Given the random selection employed in the current data, these effects were most likely washed out at the group level. However, the inclusion of twin groups in the HCP dataset affords intriguing opportunities for future exploration of individual network differences and their genetic underpinnings.

## 5. Conclusions

We have argued that network neuroscience will need to adopt methods that account for overlap between networks to fully capture the network architecture of the human brain. We have presented evidence that the mixed membership algorithm can be used to reliably generate overlapping network topographies in resting state data from multiple samples; these topographies recapitulate and extend known patterns of organization seen in nonoverlapping networks. We have argued that these overlapping networks represent a more biologically plausible organization of the brain. We have further demonstrated that regions show a broad range of multiple-network membership, and that regions that are members of networks known to be involved in cognitive control are likewise members of a wide set of networks unaccounted for in existing network models. These results highlight the need to expand our network models of cognition to fully capture the networks involved and account for the overlap and interactions between them.

## Code and data availability

A Bash- and MATLAB-based implementation of the methods used to prepare the functional connectivity matrices for assignment in this study can be accessed for open-access use at https://github.com/savannahcookson/NetChar. The mixed membership algorithm used to generate the overlapping networks used in these analyses are available open-source from https://github.com/premgopalan/svinet, courtesy of Gopalan and Blei (2013). All results derive from data that are publicly available as cited in Section 2.1 in the text.

## CRediT authorship contribution statement

Savannah L. Cookson: Conceptualization; Formal analysis; Funding acquisition; Methodology; Project administration; Resources; Software; Validation; Visualization; Roles/Writing - original draft; Writing - review & editing.

Mark D’Esposito: Conceptualization; Funding acquisition; Methodology; Project administration; Resources; Supervision; Visualization; Writing - review & editing.

## Acknowledgements

This research was supported by NIH grants 5 F32 MH119761-02 and MH63901. Data were provided in part by the Human Connectome Project, WU-Minn Consortium (Principal Investigators: David Van Essen and Kamil Ugurbil; 1U54MH091657) funded by the 16 NIH Institutes and Centers that support the NIH Blueprint for Neuroscience Research; and by the McDonnell Center for Systems Neuroscience at Washington University.

## References

1. Badre, D., & Nee, D. E. (2018). Frontal Cortex and the Hierarchical Control of Behavior. Trends in Cognitive Sciences, 22(2), 170–188. https://doi.org/10.1016/j.tics.2017.11.005

2. Baniqued, P. L., Gallen, C. L., Voss, M. W., Burzynska, A. Z., Wong, C. N., Cooke, G. E., Duffy, K., Fanning, J., Ehlers, D. K., Salerno, E. A., Aguiñaga, S., McAuley, E., Kramer, A. F., & D’Esposito, M. (2018). Brain network modularity predicts exercise-related executive function gains in older adults. Frontiers in Aging Neuroscience, 9(JAN), 1–17. https://doi.org/10.3389/fnagi.2017.00426

3. Bassett, D. S., & Bullmore, E. T. (2006). Small-world brain networks. Neuroscientist, 12(6), 512– 523. https://doi.org/10.1177/1073858406293182

4. Bijsterbosch, J. D., Beckmann, C. F., Woolrich, M. W., Smith, S. M., & Harrison, S. J. (2019). The relationship between spatial configuration and functional connectivity of brain regions revisited. ELife, 8. https://doi.org/10.7554/eLife.44890

5. Biswal, B., Yetkin, F. Z., Haughton, V. M., & Hyde, J. S. (1995). Functional connectivity in the motor cortex of resting human brain using echo-planar mri. Magnetic Resonance in Medicine, 34(4), 537–541. https://doi.org/10.1002/mrm.1910340409

6. Cocchi, L., Zalesky, A., Fornito, A., & Mattingley, J. B. (2013). Dynamic cooperation and competition between brain systems during cognitive control. Trends in Cognitive Sciences, 17(10), 493–501. https://doi.org/10.1016/j.tics.2013.08.006

7. Cohen, J. R., & D’Esposito, M. (2016). The Segregation and Integration of Distinct Brain Networks and Their Relationship to Cognition. Journal of Neuroscience, 36(48), 12083– 12094. https://doi.org/10.1523/JNEUROSCI.2965-15.2016

8. Cohen, J. R., Gallen, C. L., Jacobs, E. G., Lee, T. G., & D’Esposito, M. (2014). Quantifying the reconfiguration of intrinsic networks during working memory. PLoS ONE, 9(9). https://doi.org/10.1371/journal.pone.0106636

9. Cole, M. W., Bassett, D. S., Power, J. D., Braver, T. S., & Petersen, S. E. (2014). Intrinsic and task-evoked network architectures of the human brain. Neuron, 83(1), 238–251. https://doi.org/10.1016/j.neuron.2014.05.014

10. Cole, M. W., Reynolds, J. R., Power, J. D., Repovs, G., Anticevic, A., & Braver, T. S. (2013). Multi-task connectivity reveals flexible hubs for adaptive task control. Nature Neuroscience, 16(9). https://doi.org/10.1038/nn.3470

11. Cox, R. W. (1996). AFNI: Software for analysis and visualization of functional magnetic resonance neuroimages. Computers and Biomedical Research, 29(3), 162–173. https://doi.org/10.1006/cbmr.1996.0014

12. Dosenbach, N. U. F., Fair, D. A., Cohen, A. L., Schlaggar, B. L., & Petersen, S. E. (2008). A dual-networks architecture of top-down control. Trends in Cognitive Sciences, 12(3), 99– 105. https://doi.org/10.1016/j.tics.2008.01.001

13. Dosenbach, N. U. F., Fair, D. A., Miezin, F. M., Cohen, A. L., Wenger, K. K., Dosenbach, R. A. T., Fox, M. D., Snyder, A. Z., Vincent, J. L., Raichle, M. E., Schlaggar, B. L., & Petersen, S. E. (2007). Distinct brain networks for adaptive and stable task control in humans. Proceedings of the National Academy of Sciences of the United States of America, 104(26), 11073–11078. https://doi.org/10.1073/pnas.0704320104

14. Friston, K. J. (1994). Functional and effective connectivity in neuroimaging: A synthesis. Human Brain Mapping, 2(1–2), 56–78. https://doi.org/10.1002/hbm.460020107

15. Glasser, M. F., Coalson, T. S., Robinson, E. C., Hacker, C. D., Harwell, J., Yacoub, E., Ugurbil, K., Andersson, J., Beckmann, C. F., Jenkinson, M., Smith, S. M., & Van Essen, D. C. (2016). A multi-modal parcellation of human cerebral cortex. Nature, 536(7615), 171–178. https://doi.org/10.1038/nature18933

16. Glasser, M. F., Sotiropoulos, S. N., Wilson, J. A., Coalson, T. S., Fischl, B., Andersson, J. L., Xu, J., Jbabdi, S., Webster, M., Polimeni, J. R., Van Essen, D. C., & Jenkinson, M. (2013). The minimal preprocessing pipelines for the Human Connectome Project. NeuroImage, 80, 105–124. https://doi.org/10.1016/j.neuroimage.2013.04.127

17. Goldman-Rakic, P. S. (1988). Topography of cognition: parallel distributed networks in primate association cortex. Annual Review of Neuroscience, 11, 137–156. https://doi.org/10.1146/annurev.ne.11.030188.001033

18. Gopalan, P. K., & Blei, D. M. (2013). Efficient discovery of overlapping communities in massive networks. Proceedings of the National Academy of Sciences of the United States of America, 110(36), 14534–14539. https://doi.org/10.1073/pnas.1221839110

19. Gordon, E. M., Laumann, T. O., Gilmore, A. W., Newbold, D. J., Greene, D. J., Berg, J. J., Ortega, M., Hoyt-Drazen, C., Gratton, C., Sun, H., Hampton, J. M., Coalson, R. S., Nguyen, A. L., McDermott, K. B., Shimony, J. S., Snyder, A. Z., Schlaggar, B. L., Petersen, S. E., Nelson, S. M., & Dosenbach, N. U. F. (2017). Precision Functional Mapping of Individual Human Brains. Neuron, 95(4), 791–807.e7. https://doi.org/10.1016/j.neuron.2017.07.011

20. Gratton, C., Laumann, T. O., Nielsen, A. N., Greene, D. J., Gordon, E. M., Gilmore, A. W., Nelson, S. M., Coalson, R. S., Snyder, A. Z., Schlaggar, B. L., Dosenbach, N. U. F., & Petersen, S. E. (2018). Functional Brain Networks Are Dominated by Stable Group and Individual Factors, Not Cognitive or Daily Variation. Neuron, 98(2), 439–452.e5. https://doi.org/10.1016/j.neuron.2018.03.035

21. Honey, C. J., Sporns, O., Cammoun, L., Gigandet, X., Thiran, J. P., Meuli, R., & Hagmann, P. (2009). Predicting human resting-state functional connectivity from structural connectivity. Proceedings of the National Academy of Sciences of the United States of America, 106(6), 2035–2040. https://doi.org/10.1073/pnas.0811168106

22. Hwang, K., Bertolero, M. A., Liu, W. B., & D’Esposito, M. (2017). The Human Thalamus Is an Integrative Hub for Functional Brain Networks. The Journal of Neuroscience, 37(23), 5594– 5607. https://doi.org/10.1523/jneurosci.0067-17.2017

23. Ito, T., Kulkarni, K. R., Schultz, D. H., Mill, R. D., Chen, R. H., Solomyak, L. I., & Cole, M. W. (2017). Cognitive task information is transferred between brain regions via resting-state network topology. Nature Communications, 8(1), 1–13. https://doi.org/10.1038/s41467-017-01000-w

24. Khambhati, A. N., Mattar, M. G., Wymbs, N. F., Grafton, S. T., & Bassett, D. S. (2018). Beyond modularity: Fine-scale mechanisms and rules for brain network reconfiguration. NeuroImage, 166(September 2017), 385–399. https://doi.org/10.1016/j.neuroimage.2017.11.015

25. Kitzbichler, M. G., Henson, R. N. A., Smith, M. L., Nathan, P. J., & Bullmore, E. T. (2011). Cognitive effort drives workspace configuration of human brain functional networks. Journal of Neuroscience, 31(22), 8259–8270. https://doi.org/10.1523/JNEUROSCI.0440-11.2011

26. Langers, D. R. M. (2009). Blind source separation of fMRI data by means of factor analytic transformations. NeuroImage, 47(1), 77–87. https://doi.org/10.1016/j.neuroimage.2009.04.017

27. Menon, V. (2011). Large-scale brain networks and psychopathology: A unifying triple network model. Trends in Cognitive Sciences, 15(10), 483–506. https://doi.org/10.1016/j.tics.2011.08.003

28. Mohr, H., Wolfensteller, U., Betzel, R. F., Mišić, B., Sporns, O., Richiardi, J., & Ruge, H. (2016). Integration and segregation of large-scale brain networks during short-term task automatization. Nature Communications, 7. https://doi.org/10.1038/ncomms13217

29. Najafi, M., McMenamin, B. W., Simon, J. Z., & Pessoa, L. (2016). Overlapping communities reveal rich structure in large-scale brain networks during rest and task conditions. NeuroImage, 135, 92–106. https://doi.org/10.1016/j.neuroimage.2016.04.054

30. Nee, D. E. (2021). Integrative frontal-parietal dynamics supporting cognitive control. ELife, 10. https://doi.org/10.7554/eLife.57244

31. Parkin, B. L., Hellyer, P. J., Leech, R., & Hampshire, A. (2015). Dynamic network mechanisms of relational integration. Journal of Neuroscience, 35(20), 7660–7673. https://doi.org/10.1523/JNEUROSCI.4956-14.2015

32. Pedersen, M., Zalesky, A., Omidvarnia, A., & Jackson, G. D. (2018). Multilayer network switching rate predicts brain performance. Proceedings of the National Academy of Sciences of the United States of America, 115(52), 13376–13381. https://doi.org/10.1073/pnas.1814785115

33. Pessoa, L. (2014). Understanding brain networks and brain organization. Physics of Life Reviews, 11(3), 400–435. https://doi.org/10.1016/j.plrev.2014.03.005

34. Power, J. D., Cohen, A. L., Nelson, S. M., Wig, G. S., Barnes, K. A., Church, J. A., Vogel, A. C., Laumann, T. O., Miezin, F. M., Schlaggar, B. L., & Petersen, S. E. (2011). Functional Network Organization of the Human Brain. Neuron, 72(4), 665–678. https://doi.org/10.1016/j.neuron.2011.09.006

35. Power, J. D., Schlaggar, B. L., Lessov-Schlaggar, C. N., & Petersen, S. E. (2013). Evidence for hubs in human functional brain networks. Neuron, 79(4), 798–813. https://doi.org/10.1016/j.neuron.2013.07.035

36. Schaefer, A., Kong, R., Gordon, E. M., Laumann, T. O., Zuo, X.-N., Holmes, A. J., Eickhoff, S. B., & Yeo, B. T. T. (2018). Local-Global Parcellation of the Human Cerebral Cortex from Intrinsic Functional Connectivity MRI. Cerebral Cortex, 28(9), 3095–3114. https://doi.org/10.1093/cercor/bhx179

37. Seeley, W. W., Menon, V., Schatzberg, A. F., Keller, J., Glover, G. H., Kenna, H., Reiss, A. L., & Greicius, M. D. (2007). Dissociable intrinsic connectivity networks for salience processing and executive control. Journal of Neuroscience, 27(9), 2349–2356. https://doi.org/10.1523/JNEUROSCI.5587-06.2007

38. Seitzman, B. A., Gratton, C., Laumann, T. O., Gordon, E. M., Adeyemo, B., Dworetsky, A., Kraus, B. T., Gilmore, A. W., Berg, J. J., Ortega, M., Nguyen, A. L., Greene, D. J., McDermott, K. B., Nelson, S. M., Lessov-Schlaggar, C. N., Schlaggar, B. L., Dosenbach, N. U. F., & Petersen, S. E. (2019). Trait-like variants in human functional brain networks. Proceedings of the National Academy of Sciences of the United States of America, 116(45), 22851–22861. https://doi.org/10.1073/pnas.1902932116

39. Shinn, M., Romero-Garcia, R., Seidlitz, J., Váša, F., Vértes, P. E., & Bullmore, E. (2017). Versatility of nodal affiliation to communities. Scientific Reports, 7(1), 1–12. https://doi.org/10.1038/s41598-017-03394-5

40. Smith, V., Mitchell, D. J., & Duncan, J. (2018). Role of the default mode network in cognitive transitions. Cerebral Cortex, 28(10), 3685–3696. https://doi.org/10.1093/cercor/bhy167

41. Sporns, O. (2013). Network attributes for segregation and integration in the human brain. Current Opinion in Neurobiology, 23(2), 162–171. https://doi.org/10.1016/j.conb.2012.11.015

42. Van Essen, D. C., Smith, S. M., Barch, D. M., Behrens, T. E. J., Yacoub, E., & Ugurbil, K. (2013). The WU-Minn Human Connectome Project: An overview. NeuroImage, 80, 62–79. https://doi.org/10.1016/j.neuroimage.2013.05.041

43. Van Essen, D. C., Ugurbil, K., Auerbach, E. J., Barch, D. M., Behrens, T. E. J., Bucholz, R., Chang, A., Chen, L., Corbetta, M., Curtiss, S., Della Penna, S., Feinberg, D. A., Glasser, M. F., Harel, N., Heath, A. C., Larson-Prior, L. J., Marcus, D., Michalareas, G., Moeller, S., … Yacoub, E. (2012). The Human Connectome Project: A data acquisition perspective. NeuroImage, 62(4), 2222–2231. https://doi.org/10.1016/j.neuroimage.2012.02.018

44. Yeo, B. T. T., Krienen, F. M., & Buckner, R. L. (2014). Estimates of Segregation and Overlap of Functional Connectivity Networks in the Human Cerebral Cortex. NeuroImage, 88, 212–227. https://doi.org/10.1016/j.neuroimage.2013.10.046.Estimates

45. Yeo, B. T. T., Krienen, F. M., Sepulcre, J., Sabuncu, M. R., Lashkari, D., Hollinshead, M., Roffman, J. L., Smoller, J. W., Zollei, L., Polimeni, J. R., Fischl, B., Liu, H., & Buckner, R. L. (2011). The organization of the human cerebral cortex estimated by intrinsic functional connectivity. Journal of Neurophysiology, 106(3), 1125–1165. https://doi.org/10.1152/jn.00338.2011

